# Ovine fetuses from slaughterhouses: a useful source for neural cell primary cultures

**DOI:** 10.1101/815159

**Authors:** Vittorio Farina, Sergio Domenico Gadau, Gianluca Lepore, Marcella Carcupino, Marco Zedda

**Affiliations:** Department of Veterinary Medicine, University of Sassari, Via Vienna 2, 07100 Sassari, Italy

## Abstract

A lot of evidence demonstrates that sheep could represent an experimental model to set up medical procedures in a view of their application on humans. Sheep are chosen as model for human biomechanical studies because their skeleton has some similarities with humans. Sheep are gyrencephalic so the cerebral cortex can show valuable signposts to identify particular cortical regions, unlike rats that are lissencephalic animals. The aim of this work was to set up sheep primary cultures from ovine fetuses at different ages, from pregnant uteri retrieved at local abattoirs. Cell characterization demonstrated that one cell population was immunopositive to GFAP and identifiable as astrocytes, whereas a second cell type was III β-tubulin-positive, and hence classified as neurons. A 60 day old fetus is suitable to obtain neurons, whereas in a 90 day old fetus the cell culture is predominantly characterized by glial cells. The procedure here proposed is inexpensive. Indeed, the fetuses casually found during sheep slaughtering have no cost, unlike the classical experimental animals, such as mice, rats, rabbits, that require very high economical efforts. Finally, our protocol fully eliminates the need of animal killing, being living animals replaced by a validated in vitro model in agreement with the 3Rs statement.

## Introduction

In experimental biology, the need to turn to an experimental model which might be closer to humans is increasingly pressing. Indeed, although rodents remain the most widely used species for experimental research, other mammals may be useful because of their complex anatomy and physiology in comparison to rodents. Therefore, several papers have been published in order to underline the validity of other animal models such as dogs and cattle [1, 2, 3]. Recently, sheep tissues and organs have been used to perform experimental procedures in the studies of human pathological conditions. The scientific works carried out on sheep are very numerous and with different aims. Indeed, these animal species show similarities to humans in lots of anatomical, physiological and pathological features. By browsing the concerning literature, it can be noted that almost all anatomical systems of sheep have been considered as useful experimental models to set up medical procedures in a view of their application on humans. Sheep are chosen as model for human biomechanical studies because their skeleton has similarities with humans, for example in mineral metabolism, bone density and bone healing [4, 5, 6, 7, 8]. Sheep are also frequently used for the preclinical analyses as they represent a very useful model for the assessment of the safety and pharmacokinetics of microbicide drugs [9].

In accordance with the literature [10] there are many other advantages of using sheep as an experimental model. In neuroscience studies, sheep have acquired an increasing interest for many aspects. It is worth to note that this species is a spontaneous model for the formation of neurofibrillary tangles in the cerebral cortex [11, 12], which are typical of neurological disorders, such as Alzheimer’s disease. There are many advantages in studying the cerebral cortex of sheep as it has cerebral gyri similar to those in the human neocortex. The presence of gyri and sulci in sheep brain provides valuable signposts to identify particular cortical regions, unlike rats or mice that are lissencephalic animals [13]. Moreover, the size of the sheep brain may facilitate physiological and neuropathological comparisons with the human brain. Sections of sheep hypothalamus have been successfully studied as a model of brain sex differentiation. In particular, the morphology of the ovine sexually dimorphic nucleus, located in the central region of the preoptic area of the hypothalamus, has been examined by several authors which found interesting results applicable to many mammalian species including humans [14, 15, 16, 17,]. Cell cultures carried out from sheep brain represent another important tool for studying neural cell biology instead of the usual cell cultures obtained from rats [18, 19, 20, 21, 22]. Actually, in addition to ovine cultured cells, human-derived cell models exist, including stem cell-derived primary neuronal and glial cultures, as well as sophisticated 3D organoids [23, 24, 25]. However, such models may be expensive and/or need a high degree of manipulation in order to get differentiated neural cells. The aim of this work is to evaluate the different possibilities in setting up sheep neuronal or glial primary cultures from fetuses sampled in slaughterhouses otherwise addressed to destruction, with no animal suffering and without the use of laboratory animals.

## Material and methods

### Sampling

Fetuses at different ages were collected in local slaughterhouses (Fig.1). Sheep were healthy and regularly slaughtered since pregnancy was not known to the farmers. Indeed, during spring-summer period there are good chances to find accidentally pregnant sheep in Sardinian public slaughterhouses, as extensive farming is conducted in Sardinia and therefore sheep are in constant presence of the ram. In that period, about 10% of sheep may be found casually pregnant. The sampling took place on dead sheep fetuses contained in the gravid uteri destined for disposal, so that neither anesthesia nor euthanasia were performed for the present study. All good practices on animal welfare have been adopted by the slaughterhouse.

**Fig. 1.**
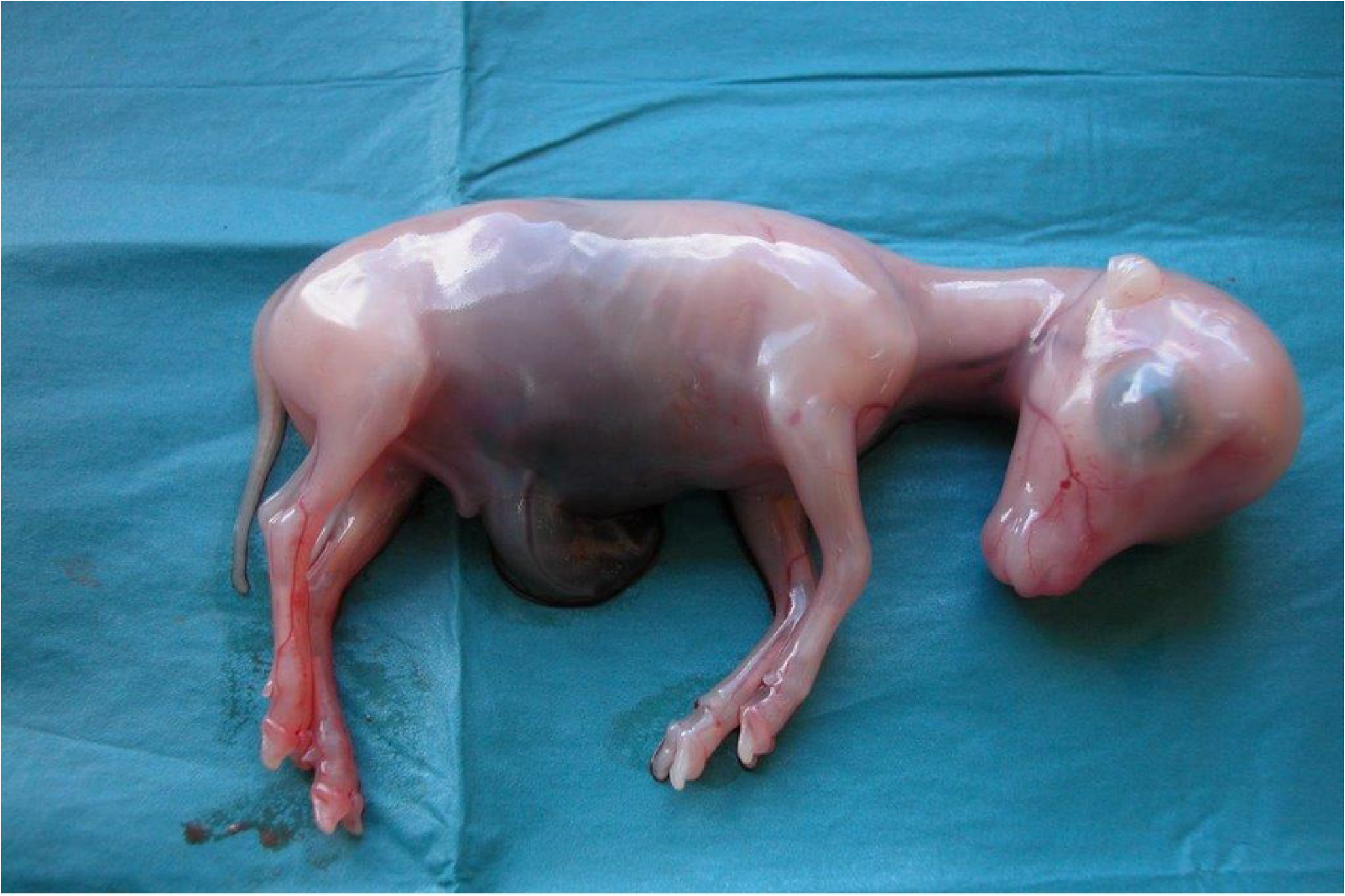
Ovine embryo at 40 gestation days. The period of gestation was estimated on the basis of the fetal crown-rump length.

Within half an hour the fetuses were transported to our laboratory at 39°C, which is the basal temperature of this species [26]. First of all, the period of gestation was estimated on the basis of the fetal crown-rump length following the tables reported in the literature [27]. Based on these measurements, fetuses at three different gestation ages (40, 60, 90 days) were collected.

Five brains from each gestational period were collected, for a total of fifteen brains. Homogenates were mechanically obtained (Turrax, IKA, Staufen, Germany) from each whole fetal brain and then suspended in an ice-cold freezing medium containing Dulbecco’s Modified Eagle’s Medium (DMEM), 1% HEPES (pH 7.4), 1% fetal calf serum and 10% dimethylsulfoxide (DMSO). Tissues were then stored in liquid nitrogen until use.

### Cell culture procedures

Frozen tissues were rapidly thawed in a water bath at 39°C and minced in small fragments under sterile conditions into a laminar flow hood. A papain dissociation kit (Worthington Biochemical Corporation, Lakewood, NJ, USA) was used to dissociate cells following the guidelines of the Company. Cells were then suspended in a basal medium consisting of 1:1 DMEM and Ham’s F12, supplemented with penicillin (30mg/l), streptomycin (50mg/l), sodium bicarbonate (2.4g/l), insulin (10µg/ml), transferrin (10µg/ml), sodium selenite (10^−8^M) and 10% fetal calf serum, at 39°C in a 5% CO_2_ humidified atmosphere. Cells were plated at the rate of 5×10^5^ on glass coverslips, previously coated with poly-L-lysine. All reagents, when not specified, were from Merck, (Darmstadt, Germany).

### Immunofluorescence staining

In order to distinguish neurons and astrocytes, a double immunofluorescence method was performed on the monolayers obtained at the scheduled times. At day 10 of culture cells were fixed with 4% paraformaldehyde at room temperature for 30min, washed with PBS, and then permeabilized with 0.1% Triton X-100 for 15min at room temperature. Subsequently they were incubated overnight with anti-class III β-tubulin antibody (monoclonal, clone DM1A), a widely-recognized marker of neuronal cells [28] diluted 1:200, and then for 1h at 37°C with anti-mouse FITC-conjugated antibody diluted 1:100 (AlexaFluor 488, Termo Fisher, Waltham, MA, USA). Cells were then incubated for 1h with polyclonal anti-GFAP antibody, diluted 1:200, specific for glial fibrillary acidic protein present in astrocytes. Finally, an incubation of 1h with a secondary anti-rabbit TRITC-conjugated antibody, diluted 1:100 (AlexaFluor 594) was carried out. In addition, nuclei were counterstained with Hoechst #33258 fluorescent dye. Additional monolayers were used as negative control by omitting the primary antibody or employing non-immune mouse serum. Cells were finally observed with a Leica TCS SP5 DMI 6000CS confocal microscope (Leica Microsystems GmbH, Wetzlar, Germany). FITC was excited at 488nm and emission was detected between 510 and 550nm whereas TRITC was excited at 568nm and emission was detected between 585 and 640nm. Microscopic pictures of monolayers were taken, each including more than 50 cells. In order to evaluate the number of neurons and astrocytes, we proceeded as follows: five high-magnification photos acquired by confocal microscope were taken for each monolayer (15 in total). A grid composed of twelve squares (387µm^2^) was placed on each picture using ImageJ freeware analysis software. Through the Analyze/Counter plugin, we proceeded to count the cells which were positive to anti-β III tubulin antibody (neurons) and then those positive to anti-GFAP (astrocytes). Cell number was expressed in percentage. The counting was repeated by three different researchers to evaluate possible inter-observer differences which were lower than 2%.

### Statistics

All data was expressed as means ±SD and tested by One-Way ANOVA. Statistical analyses were performed by using Statigraphics software (Statgraphics Technologies, Inc., The Plains,VA, USA) and a *p*< 0.05 was considered significant.

## Results

Technical procedures led to setting up neural primary cultures starting from all the gestation periods taken into consideration (40^th^, 60^th^ and 90^th^ day), obtaining numerous and viable cells. At gestational days 60 and 90, cells were 40% and 110% more numerous than at gestational day 40 respectively (Fig. 2). Cell plates from the five brain fragments belonging to the same gestational period were homogenous as to morphology and growth rate after 10 days of cultures.

**Fig. 2.**
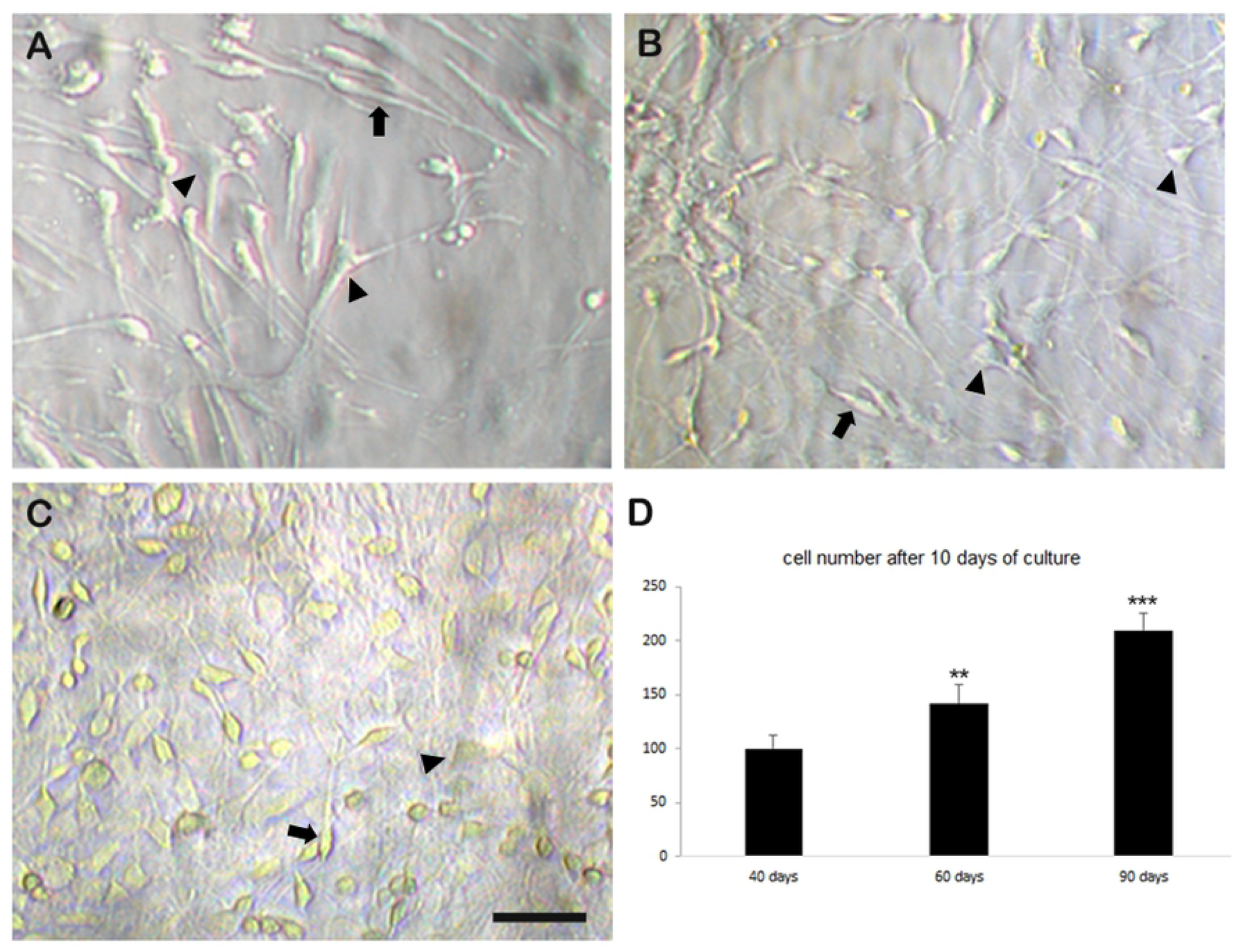
Phase contrast optics. Evaluations of primary cell cultures from fetal sheep brain at different development stages: 40 (A), 60 (B) and 90 (C) gestation days. Arrows indicate spindle shaped cells; arrowheads indicate stellate cells. Bar=60 µm. The diagram (D) shows the different amount of cells after 10 days of culture. Data are expressed in percentage in comparison to the value of cell numbers from 40 day old fetuses. ** *p*<0.01, \*\*\**p*<0.001.

Two cell types were mostly present: i) stellate cells showing a pleomorphic shape, large flat cell bodies and cytoplasmic processes, ii) cells with spindle-like or triangular-shaped cell bodies, with long processes that tended to have contacts with those of contiguous cells. The immunofluorescence characterization of these two cell populations testified that the first cell-type was immunopositive to GFAP and thus identifiable as astrocytes, whereas the second type was formed by neurons as they were III β-tubulin-positive (Fig. 3). Although both cell types were present at all the three scheduled times, the two populations showed different percentages as reported in Fig. 3. Specifically, at the 40^th^ day of pregnancy cultured cells were prevalently neurons (60%) and at the 60^th^ day, neurons were even more numerous (90%, *p*<0.0001). On the other hand, at the 90^th^ day the most numerous cells were astrocytes (80%, *p*<0.001). The differences in the number of astrocytes at 40 and 60 days and the number of neurons at 90 days was not statistically significant.

**Fig. 3.**
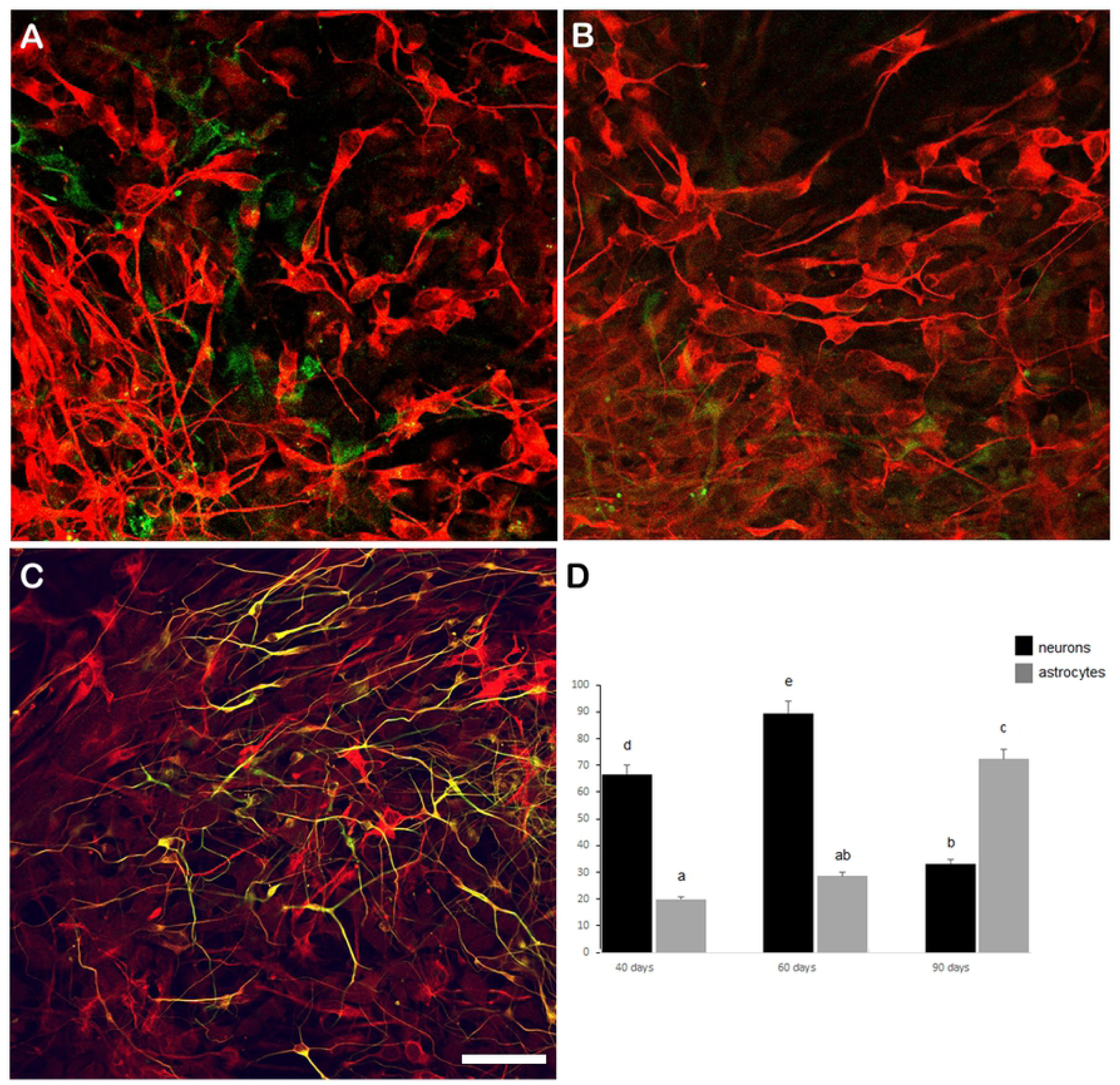
Double immunofluorescence staining. This method shows the two different cell populations: immunopositive neurons to anti-β III tubulin (red), and immunopositive astrocytes to anti-GFAP (green) on cultures belonging to fetal ovine brain explants of 40 (A), 60 (B) and 90 (C) days of gestation. Bar = 25 µm. The diagram (D) highlights the percentage of neurons and astrocytes at different stages of fetal development. ANOVA One Way. Lower case letters indicate statistical differences (c, d, e, *p*<0.0001).

## Discussion

Primary neural cell cultures represent a useful protocol to study cell morphology, cell physiology and toxicology [29,30]. Recently, the research of experimental models suitable to humans has increased. Indeed, the favorite animal models are those showing brain size, brain complexity, life span expectancy and diseases similar to those of humans [31, 32]. Human-derived cell cultures exist, including stem cell-derived primary neuronal and glial cultures, as well as sophisticated 3D organoids [23,24,25]. However, cultures from lots of animal species are also exploited as validated models. In this work, ovine fetuses at different developmental stages have been employed in order to obtain neural primary cell cultures, so that a comparison could be possible to cultures set up from traditional laboratory animals. Our results show that sheep primary neural cell cultures are very useful as they allow to obtain significant amounts of viable neuronal and glial cells in an easy manner. Furthermore, our work confirms the usefulness of the dissociation method with papain. Such enzyme allows a mild dissociation, giving the possibility to obtain a high number of very viable neural cells [33, 34]. Moreover, as already highlighted [1], our experiment confirms the usefulness of the cryopreservation of explants, as it allows to obtain a proliferating culture even after years from the collection. Concerning the different gestational periods of fetuses from which the samples were taken, the data obtained demonstrates to be original. It is known that in rodents fetal age may affect the amount of neural cells obtainable [35, 36, 37] and our data provides information on ovine species for the first time. From the data obtained in this work, we can assert that cell collection at the 60^th^ day of pregnancy is optimal if researchers wish to set up a primary culture where neurons are prevalent, whereas cell collection at the 90^th^ day of pregnancy is suitable if a culture of astrocytes is required. Sometimes a co-culture of neurons and astrocytes is needed, for example when the research is focused on the relationships between the two neural cell types. In this case, sheep fetuses represent a useful source of brain tissue, since a co-culture of neurons and astrocytes can be easily obtained at the 40^th^ day of pregnancy. Unlike other traditional laboratory animals, sheep fetuses in our protocol came from ewes intended to be slaughtered and hence there are no scheduled sacrifices of living animals.

This procedure avoids euthanasia, in accordance to the guidelines of the 3Rs Program [7]. In the last decades, alternatives to animal testing have been proposed in order to enforce the strategy of 3Rs principles (Reduction, Refinement and Replacement of laboratory use of animals) [38, 39, 40]. Despite their large use, the experimental models based on mice and rats show some limitations when compared to the investigations of large mammals [41], such as the inbred nature of certain rodent strains and a short life span. Therefore, various Authors have proposed several experimental models of animal species closer to humans than mice and rats, such as dogs, cows and sheep [42, 2]. Currently the general trend towards saving animal lives in experimental studies is increasing, and animal welfare is becoming of primary importance at all experimental stages. The replacement of animals with a validated alternative method, such as *in vitro* culture, is suggested. The present protocol is consistent to the principle that the experimental model must be carefully chosen to ensure the analogy to humans, to avoid excessive use of animals and to prevent waste of time, effort and high costs [5].

## Conclusions

The *in vitro* procedure here described is inexpensive. On the other hand the use of experimental animals such as mice, rats, etc., requires very high economical efforts, since they must be housed in suitable enclosures according to the law. The collection at the slaughterhouse followed by an adequate cryopreservation in liquid nitrogen of samples otherwise destined for destruction allows the availability of samples for the preparation of long lasting cell cultures. Finally, our protocol fully eliminates the need of animal killing, being living animals replaced by a validate in vitro model.

## Acknowledgements

The Authors wish to thank Dr. Nicolò Giordano for his peer review of the English manuscript.

